# A mass spectrometry-based atlas of extracellular matrix proteins across 25 mouse organs

**DOI:** 10.1101/2022.03.04.482898

**Authors:** Maxwell C. McCabe, Anthony J. Saviola, Kirk C. Hansen

## Abstract

The extracellular matrix is a critical non-cellular component of multi-cellular organisms containing a variety of proteins, glycoproteins, and proteoglycans which has been implicated in a wide variety of essential biological processes, including development, wound healing, and aging. Due to low solubility, many ECM proteins have been underrepresented in previous proteomics datasets. Using an optimized 3-step decellularization and ECM extraction method involving chaotrope extraction and digestion via hydroxylamine hydrochloride, we have generated coverage of the matrisome across 25 organs. We observe that the top 100 most abundant proteins from the ECM fractions of all tissues are generally present in all tissues, indicating that tissue matrices are principally composed of a shared set of ECM proteins. However, these proteins vary up to 4,000-fold between tissues, resulting in highly unique matrix profiles even with the same primary set of proteins. A data reduction approach was used to reveal related networks of expressed ECM proteins across varying tissues, including basement membrane and collagen subtypes.

## Introduction

The extracellular matrix is a non-cellular network containing a variety of proteins, such as collagens, fibronectins, and laminins, as well as additional glycoproteins and proteoglycans. This three-dimensional network provides structural support to tissues, while interactions between ECM components and cell surface receptors transduce signals involved in cell migration, differentiation, and growth^1–3^. At a higher level, the modulation of ECM properties and signaling has been implicated in a wide variety of essential biological and pathological processes^4^, including wound healing^5,6^, tumor proliferation^7^, and fibrosis^8^.

Preliminary draft proteomes published in the past several years^9–11^ have demonstrated that the calculated abundance of numerous ECM proteins (e.g. collagens I, IV, V, elastin, etc.) is lower than expected^12^. This discrepancy is likely due to the high percentage (up to 85%) of the fibrillar ECM residing in a chaotrope resistant insoluble ECM fraction (iECM)^13^. In the past 15 years, there has been a large focus on generating protein atlases to describe the composition of various tissues in drosophila, mice, and humans. However, characterization of the iECM via standard proteomic methods is limited due to difficulties associated with protein solubility, thus it has not yet been fully characterized by previous tissue proteomic studies.

Despite the vast amount of data available in ProteomicsDB^14^, a repository of human proteomics datasets from more than 78 studies, collagen and other insoluble ECM proteins are underrepresented. For example, the skin dataset shows that albumin (ALB) levels are more than 4 orders of magnitude higher than collagen type 1 alpha 1 (COL1A1), and there is no proteomic evidence for elastin, which is known to be rich in the dermis^15^. Another protein atlas, PAXDb^16^, aggregates proteomic data for a variety of organisms. Across all *Mus musculus* organs, COL1A1 is ranked as the 1807^th^ most abundant protein across all organs studied, again likely underrepresenting its ubiquitously reported high abundance. A deeper, more comprehensive analysis of ECM composition across various organs and organisms is necessary to aid in both our understanding of normal and disease pathologies, aging and optimization of ECM scaffolds used in cell culture models and regenerative medicine strategies.

Two-dimensional (2D) cell culture has been the predominant technique for analysis of cellular function and fundamental biomolecular mechanisms since the mid-20^th^ century. However, within tissues dynamic reciprocity between the cell and the surrounding matrix directly influences cell behavior, causing 2D cell culture to inaccurately represent relevant *in vivo* biological conditions^17^. In the absence of physical and biochemical cues from the matrix, cell morphology, proliferation, and cell surface receptor expression are frequently altered, resulting in experimental outcomes with reduced *in vivo* correlation^18^. Three-dimensional (3D) cell culture is able to address some of these issues by providing structural and biochemical cues which better replicate the *in vivo* tissue microenvironment^18^. Since its initial development, 3D cell culture has been effectively used in many contexts to culture organoids from uniform stem cell populations which recapitulate functional features of the native tissue^19–21^. These organoids provide a novel way to model complex cellular structures that more closely resemble *in vivo* biology without the use of an entire organism^22^. Likewise, it becomes feasible to study human systems when animal models do not fully recapitulate a specific physiology or disease state.

Although engineered matrices commonly used in 3D cell culture have the advantage of easily defined and tunable composition^23^, they do not fully represent the range of biomolecules present in native ECM which direct cellular behavior. Within organotypic 3D cell culture, native matrices derived from relevant tissue sources frequently outperform more general-purpose ECM scaffolds such as purified Collagen Type I and Matrigel^24,25^. This discrepancy highlights the need to better understand the ECM composition between tissues in order to identify components which positively contribute to tissue- and organ-specific cell development.

We have developed optimized protocols for extraction of ECM that have been refined using samples from cell culture^26^, mouse tumors^27^, over a dozen mouse organs^28,29^, and extinct mammal tissues^30^, allowing for molecular-scale assessment of tissue ECM proteome composition. By performing compartment-resolved proteomic analysis on various tissue components using this ECM extraction method, we aim to characterize the composition and relative abundance of ECM protein components across 25 organs, working toward the generation of a tissue ECM proteome database.

## Methods

### Sample preparation for LC-MS/MS

Tissue samples were processed as previously described^31^. Briefly, fresh frozen samples were milled in liquid nitrogen with a mortar and pestle before lyophilization. Five milligrams of samples were processed by a stepwise extraction with CHAPS and high salt, guanidine hydrochloride and chemical digestion with hydroxylamine hydrochloride (HA) in Gnd-HCl resulting in cellular, soluble ECM (sECM), and insoluble ECM (iECM) fractions for each sample, respectively. Approximately 100 mg of 3mm glass beads were used to mechanically agitate samples in a Bullet Blender (NextAdvance) prior to all cellular and sECM extraction steps. Protein concentration of each fraction for each sample was measured using A660 Protein Assay (Pierce™). Proteolytic digestion was carried out according to the FASP protocol^32^ with 10 kDa molecular weight cutoff filters (Sartorius Vivacon 500 #VN01H02) using 30 ug of protein resulting from each fraction. Samples were prepared by reducing, alkylating, and digesting with trypsin (1:100) at 37ºC for 14 Hrs. Peptides were recovered from the filter using successive washes with of 50 mM ammonium bicarbonate and 0.1% formic acid. Final volume was adjusted to inject 6 µgs of protein.

### LC-MS/MS analysis

Global proteomics for all comparative method testing was carried out (n=3 per group) on a Fusion Lumos Tribrid mass spectrometer (Thermo Fisher Scientific) coupled to an EASY-nLC 1200 (Thermo Fisher Scientific) through a nanoelectrospray LC − MS interface. Eight μL of each sample was injected into a 20 μL loop using the autosampler. The analytical column was then switched on-line at 400 nl/min over an in house-made 100 μm i.d. × 150 mm fused silica capillary packed with 2.7 μm CORTECS C18 resin (Waters; Milford, MA). LC mobile phase solvents consisted of 0.1% formic acid in water (Buffer A) and 0.1% formic acid in 80% acetonitrile (Buffer B, Optima™ LC/MS, Fisher Scientific, Pittsburgh, PA). After 22 uL of sample loading at a maximum column pressure of 700 bar, each sample was separated on a 120-min gradient at a constant flow rate of 400 nL/min. The separation gradient for cell fractions consisted of 6% buffer B from 0 to 3 minutes, followed by a linear gradient from 6 to 42% buffer B from 3 minutes to 105 minutes. Linear gradients from 6 to 36% and 6 to 24% buffer B were utilized from 3 to 105 minutes for the sECM and iECM fractions, respectively. Gradient elution was followed by a linear increase to 55% buffer B from 105 to 110 minutes and further to 95% buffer B from 110 to 111 minutes. Flow at 95% buffer B was maintained from 111 minutes to 120 minutes to remove remaining peptides. Data acquisition was performed using the instrument supplied Xcalibur™ (version 4.5) software. The mass spectrometer was operated in the positive ion mode. Each survey scan of m/z 375–1600 was followed by higher energy collisional dissociation (HCD) MS/MS (30% collision energy) using the standard AGC target and a 35 ms maximum injection time with an isolation width of 1.6 m/z. The orbitrap was used for MS1 and MS2 detection at resolutions of 120,000 and 50,000, respectively. Dynamic exclusion was performed after fragmenting a precursor 1 time for a duration of 45 sec. Singly charged ions were excluded from HCD selection.

### Data processing

Raw data was searched using an in-house Mascot server (Version 2.5, Matrix Science) through Proteome Discoverer (ThermoFisher, version 2.5). Precursor tolerance was set to ±10 ppm and fragment tolerance was set to ±0.04 Da, allowing for 2 missed cleavages. Data was searched against SwissProt (17,029 sequences) restricted to *Mus musculus* using version 1.1 of the CRAPome for common contaminants^33^. Trypsin specific cleavage was used in searches for cell and sECM fractions, while HA/Trypsin specificity was used for hydroxylamine digested fractions. HA/Trypsin specificity was defined as cleaving C-terminal of K and R but not before P, as well as C-terminal of N but not before C, F, H, I, M, N, Q, S, W, or Y based on previous HA cleavage data^31^. Fixed modifications were set as carbamidomethyl (C). Variable modifications were set as oxidation (M), oxidation (P) (hydroxyproline), Gln->pyro-Glu (N-term Q), deamidated (NQ), and acetyl (Protein N-term). Results were filtered to 1% FDR at the peptide level using the Percolator node and at the protein level using the Protein FDR Validator node. LFQ was performed using the Precursor Ions Quantifier node, including unique and razor peptides. Data was normalized to total LFQ intensity by fraction, so that total signal in each fraction was equal across all tissues before fractions were aggregated for analysis. The Matrisome DB was used to annotate the core matrisome (ECM proteins) and matrisome associated proteins here^34^.

### Data analysis

PHATE Clustering^35^: Only data corresponding to matrisome peptides and proteins was used for PHATE clustering. Peptide intensities were first aggregated with summed protein intensities in a single data input. Data was filtered so that no entry (peptide or proteins) contained more than 10 missing values and each entry contained >5×10^7^ total intensity across all samples. Filtered data was normalized using the library.size.normalize function (*phateR* v1.0.7) and square root transformed before clustering. PHATE clustering was performed using *phateR* version 1.0.7 with default settings and the initial seed set to 2. Plotting was performed using *plotly* version 4.9.3. K-means Clustering: K-means clustering was performed using the *stats* package version 4.0.3 in R. Cluster number was optimized using *factoextra* version 1.0.7 and set at 7. K-means clustering was performed using the Lloyd algorithm with nstart and inter.max set to 100. All other settings were default. Heatmap generation: Solubility heatmaps were generated using *heatmaply* version 1.2.1 in R with default settings. Top protein heatmap (Figure 2C) was generated using Morpheus (Broad Institute, https://software.broadinstitute.org/morpheus/). Principal component analysis (PCA): PCA was performed using the *factoextra* R package version 1.0.7 with intensity data for all matrisome proteins. Data was log transformed before analysis. Data was 0 centered but not scaled within the prcomp function.

## Results

To produce baseline characterization of ECM composition across a range of different tissue types, dissected mouse organs were cryo-milled and lyophilized before being decellularized using CHAPS/NaCl buffer. The resulting ECM-enriched pellet was processed via guanidine hydrochloride (Gnd-HCl) extraction to produce a soluble ECM fraction (sECM) followed by hydroxylamine hydrochloride (HA) digestion, to produce insoluble ECM (iECM) fraction as previously described^31^ (Figure 1A). Global proteomics was performed on all extracts, resulting in 8559 high-confidence protein identifications with an average of 6952 proteins per organ. Of these identifications, 432 were assigned to matrisome proteins (Figure 1A).

**Figure 1.**
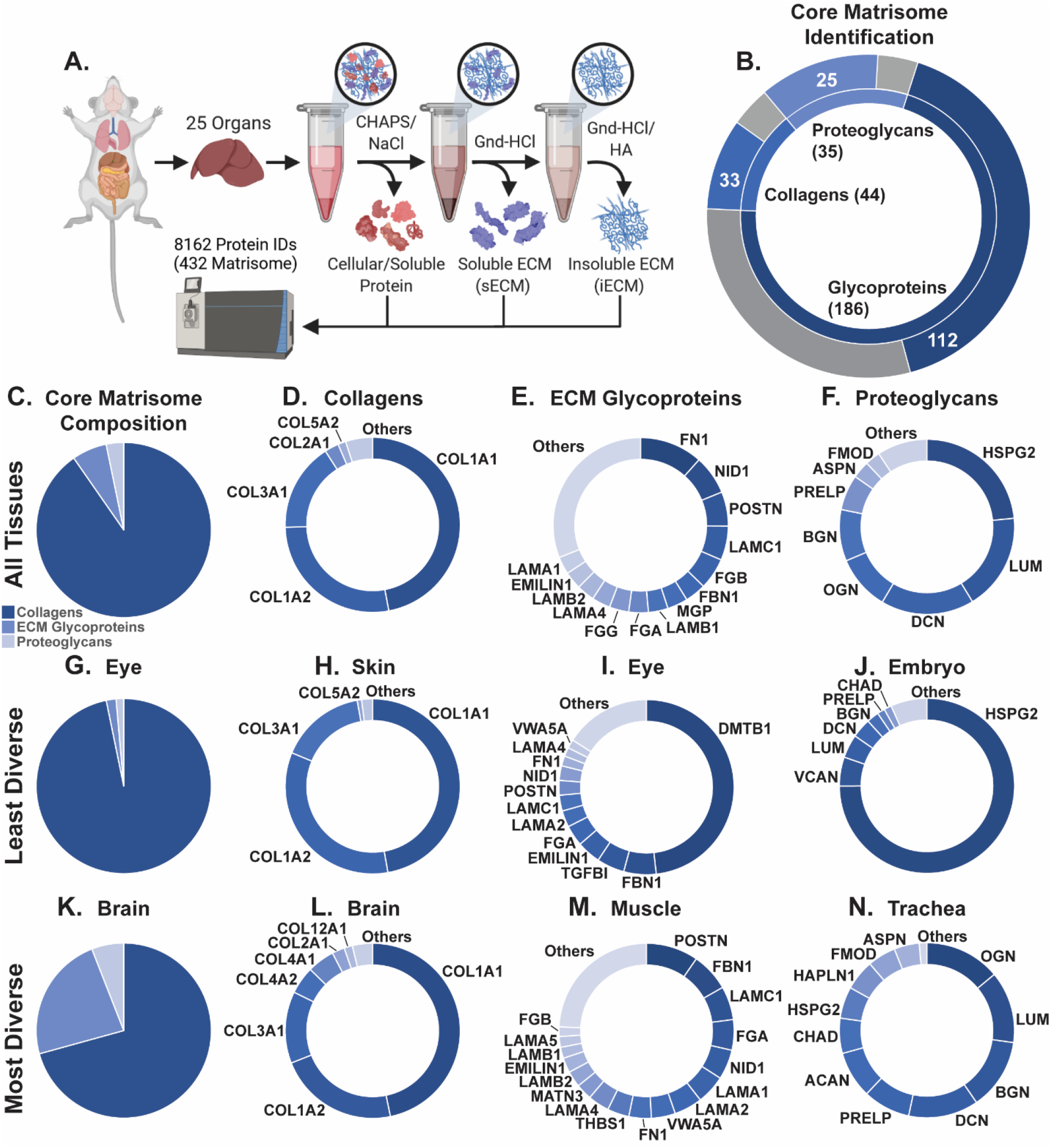
Workflow overview for sample preparation and MS analysis. A) Isolated mouse organs (27 organs, 12 with N=3) were cryo-milled, lyophilized, and delipidated. Milled samples were extracted via our previously published decellularization and ECM extraction method. B) Core matrisome proteins identified out of total from each core matrisome category. Numbers in center of plot represent total proteins in each matrisome category. Blue regions and white numbers indicate identified proteins, while grey regions indicate proteins present in MatrisomeDB but not identified. C) Percent of core matrisome signal assigned to each protein category. D-E) Top collagens (D), ECM glycoproteins (E), and proteoglycans (F) by total signal across all analyzed tissues. G-J) Core matrisome composition and top core matrisome proteins for the least diverse tissue by category. K-N) Core matrisome composition and top core matrisome proteins for the most diverse tissue by category.

Using a 3-step, ECM-optimized extraction on 25 organs, we identified 170 out of 274 theoretical matrisome proteins across all tissues, including 33 out of 44 collagens, 25 out of 35 proteoglycans, and 112 out of 186 ECM glycoproteins (Figure 1B). Core matrisome signal was composed of 90.3% collagens, 6.4% ECM glycoproteins, and 3.2% proteoglycans. Collagen composition is less diverse than other core matrisome categories, with COL1A1, COL1A2, and COL3A1 making up more than 90% of the total collagen signal across all 25 organs (Figure 1C). On the other hand, the top 3 ECM glycoproteins (FN1, NID1, and POSTN) across all samples make up only 25.1% of the total signal, with other highly abundant glycoproteins making up the remaining 75% of the category (Figure 1D). HSPG2, LUM, and DCN represent the most abundant proteoglycans across all tissues with 58.5% of the total proteoglycan signal (Figure 1E).

ECM diversity, defined as the number of proteins which compose the top 90% of signal from each core matrisome category, varies greatly between analyzed organs. Figure 1G-N displays the most and least diverse organs with respect to each core matrisome category, highlighting these differences. In skin, the least diverse organ in terms of collagen composition, collagen I makes up more than 80% of collagen signal alone (Figure 1H). In brain, on the other hand, we observe only 69% of collagen signal assigned to collagen I, with COL3A1, COL4A1, and COL4A2 also included in the top 90% of collagen signal (Figure 1L). While DMTB1 accounts for more than 48% of the ECM glycoprotein signal in the eye, the highest abundance ECM glycoprotein in muscle (POSTN) accounts for only 10% of signal for that category (Figure 1I,M). Even more drastic differences in diversity are observed for proteoglycans, where the top protein (HSPG2) accounts for 75% of proteoglycan signal in the embryo while OGN composes only 14% of signal in the trachea (Figure 1J,N).

Principal component analysis reveals clear separation between organs based on ECM composition (Figure 2A). Within the upper-right PCA quadrant, tissues (and a limb) with closely related ECM composition include leg, eye, muscle, skin, and femur. In all of these tissues matrisome signal is >85% collagen with a smaller proportion of non-structural ECM components than other tissues. Of the 168 identified core matrisome proteins in this study, 95 (56.5%) were shared between all tissues, while only 2 (NTN1 and VWA2) were unique to a single tissue sample (Figure 2B). This contrasts with a recent review of primarily cellular proteomic datasets across 8 mouse organs, which found that 46.9% of identified protein groups were unique to a single organ and only 0.16% of protein groups were identified in all organs^36^. For comparison, 44% of identified proteins were identified in all tissues within the dataset generated by the Human Protein Atlas (HUPA) project^37^. In general, we find that differences in tissue ECM composition are driven by changes in protein abundance up to 4 orders of magnitude rather than the binary presence or absence of ECM proteins.

**Figure 2.**
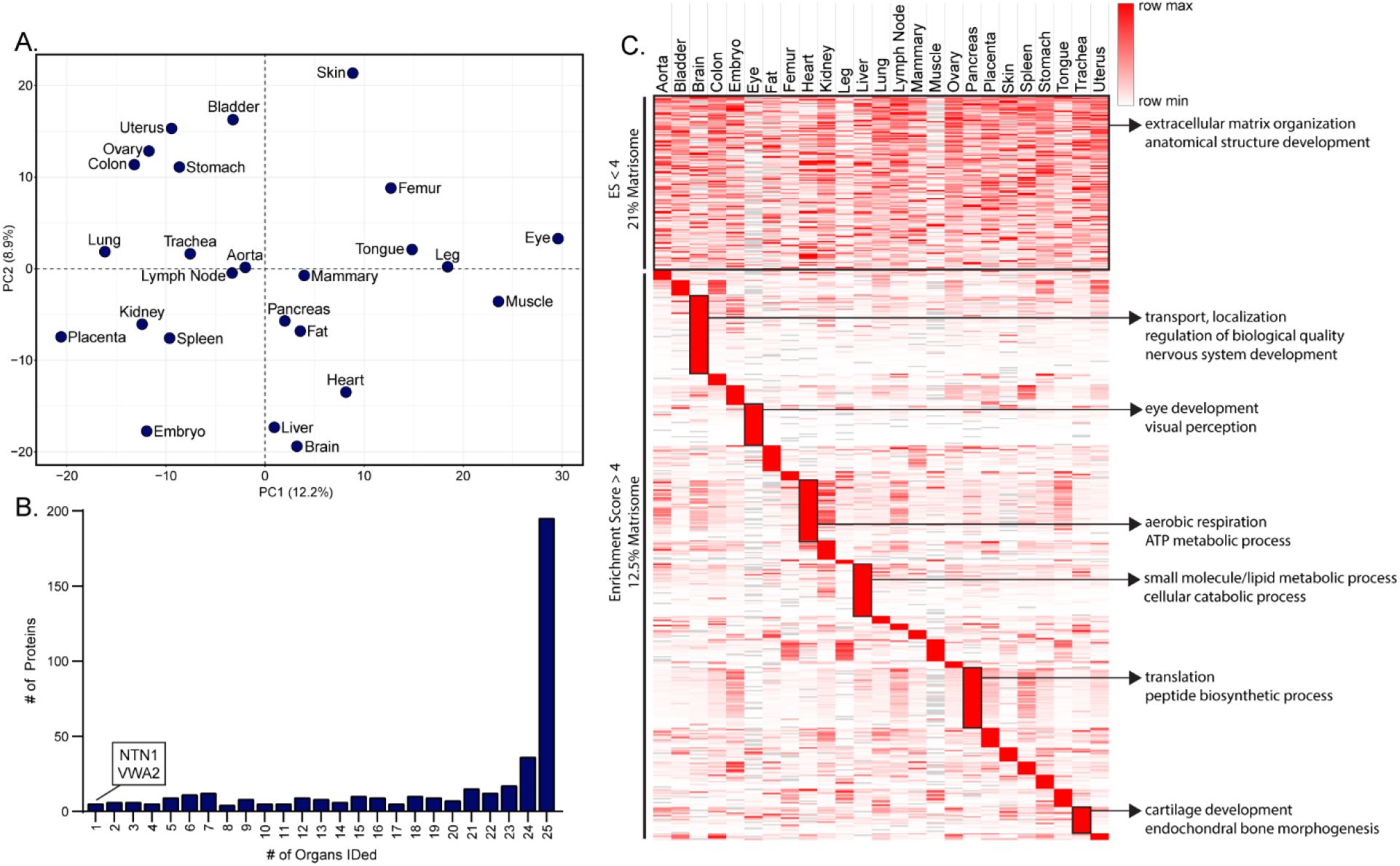
Tissues are composed of both shared structural ECM and tissue-enriched components. A) Principal component analysis (PCA) of tissue composition based on matrisome proteins. B) Number of matrisome proteins shared between tissue samples. C) Heatmap containing the top 100 proteins by intensity from the ECM fractions of all analyzed tissues. Enrichment score was calculated as the maximum intensity value for each protein divided by the mean intensity across all tissues.

Protein enrichment scores (ES) were calculated as the maximum intensity value for each protein divided by the mean intensity across all tissues. Analysis of the top 100 most abundant proteins from the ECM fractions of each organ revealed a cluster of proteins with low enrichment across tissues (ES < 4) containing 23.2% of the top 100 ECM-fraction proteins from all tissues, as well as protein clusters which are enriched in one or more tissues (ES > 4) containing the remaining 76.8% (Figure 2C). In the HUPA database generated via antibody staining, 40.9% of identified proteins are denoted as low enrichment (ES<4)^37^. The higher proportion of shared proteins in this dataset is likely due to issues with dynamic range in antibody-based quantification, resulting in an underestimate of tissue-specific enrichment.

Over 95% of the top 100 ECM-fraction proteins are present in every analyzed organ, indicating that, while often highly enriched, the most abundant proteins within the ECM fractions are generally not unique to a specific tissue. Within the low-enrichment protein cluster, 21% of proteins were assigned to the matrisome (13.5% core matrisome), compared to only 12.5% of proteins within the high enrichment group (9.3% core matrisome) (Figure 2C). Additionally, GO-term analysis reveals overrepresentation of proteins annotated as “extracellular matrix organization” and “anatomical structure development” within the low-enrichment group. The overrepresentation of matrisome proteins within the low-enrichment cluster suggests that matrisome proteins, and especially core matrisome proteins, generally vary less between tissues than cellular components. We observe strong enrichment of functionally specific GO-terms in each organ, including aerobic respiration in the heart, metabolism in the liver, and cartilage development in the trachea (Figure 2C).

By calculating the proportion of signal for each protein that is identified in the final iECM fraction of the extraction, a measure of resistance to extraction is obtained for ECM proteins across tissues. Analysis of ECM protein solubility highlights the need for additional digestion after chaotrope extraction to achieve full ECM coverage. Some core ECM components are entirely soluble in HS/CHAPS or 6M Gnd-HCl buffers, including VCAN, BCAN, and FMOD (Figure 3).

**Figure 3.**
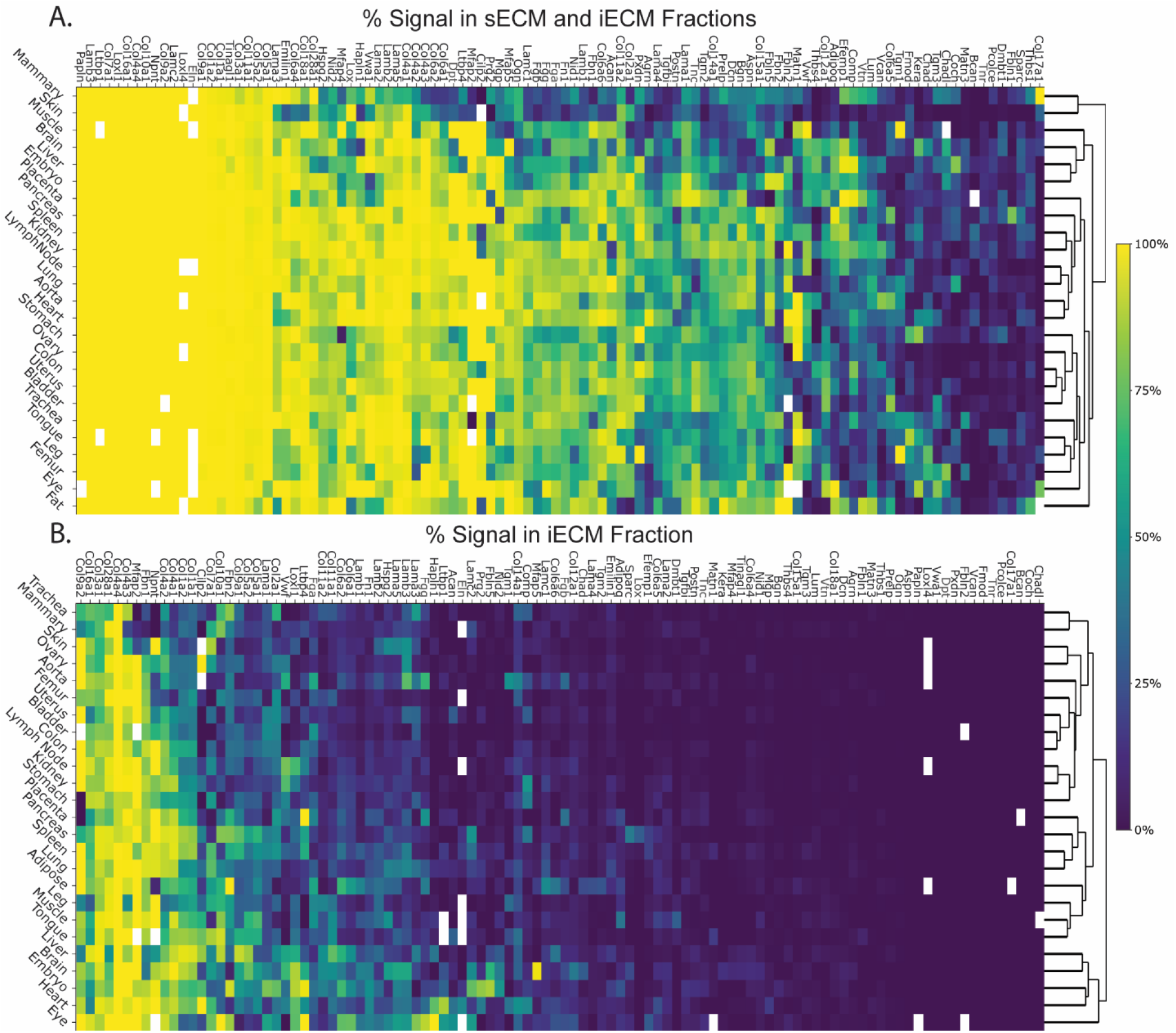
ECM protein solubility varies by protein and tissue. Cells are colored by solubility, calculated as the percent of total intensity for each protein identified in both the sECM and iECM fractions (A) and in the iECM fraction alone (B).

Other core ECM proteins, including COL3A1, COL4A3, COL4A4, and MFAP2, are highly resistant to chaotrope extraction, resulting in greater than 75% of the signal for these proteins being identified in the final iECM fraction (Figure 3B). Therefore, without chemical digestion of the remaining insoluble pellet after chaotrope extraction, these proteins remain insoluble and ECM compositional readouts are biased as a result. The effectiveness of protease digestion in a chaotrope has been evaluated and also does not access a significant fraction of this iECM^38^. Additionally, we observe some highly insoluble cellular proteins with >75% of signal in the insoluble fraction, including CASC3, LSR, and SEC16A, indicating that additional digestion of insoluble material can aid in coverage of specific cellular proteins as well as ECM (Supplementary Figure 2).

Protein solubility has previously been shown to change with aging^39^ and disease^40^. Here, we show that solubility of ECM proteins also drastically varies between tissues, with the chaotrope-insoluble percentage of highly variable proteins such as FBN1 varying as much at 20-fold between tissues (Figure 3). Other structural ECM proteins, such as COL10A1, range from being entirely chaotrope soluble in tissues such as the stomach (2.9% insoluble) to requiring chemical digestion for identification in the eye (100% insoluble). Additionally, some ECM modifiers display similar behavior, such as LOXL1, which varies from 89.1% chaotrope insoluble in embryo to 1.0% insoluble in trachea.

The ability to measure resistance to extraction across the 3 fractions as a surrogate for protein solubility represents a significant benefit of using a 3-step extraction approach. The varying resistance of ECM proteins to chaotrope extraction underscores the need for additional digestion steps to access structural ECM components. In addition, digestion of the insoluble fraction can lead to the detection of highly insoluble cellular proteins that might not be identified using alternative extraction methods.

Utilizing subcategory annotations applied to the contents of MatrisomeDB^34^ (supplementary data), we compared the relative abundance of protein subclasses across tissues. In all organs, fibrillar collagen composes >85% of the total collagen signal, with brain containing both the lowest proportion of fibrillar collagen and the lowest amount of collagen overall (Figure 4A,B). While COL1A1 is the most abundant fibrillar collagen across all tissues, the percent of signal assigned to lower-level fibrillar collagens varies between samples. Collagen 3 makes up a highly variable percentage of total collagen signal across tissues, composing 34.1% of pancreas collagen but only 2.6% of collagen in the femur. When comparing the fractional composition of different classes of non-fibrillar collagens across tissues, drastic variation in the most prevalent collagen subtype was observed. For example, while both placenta and uterus contain similar amounts of non-fibrillar collagens in total, network forming collagens represent 58% of the total non-fibrillar collagen signal in placenta, but only 7% of the signal in uterus (Figure 4C,D).

**Figure 4.**
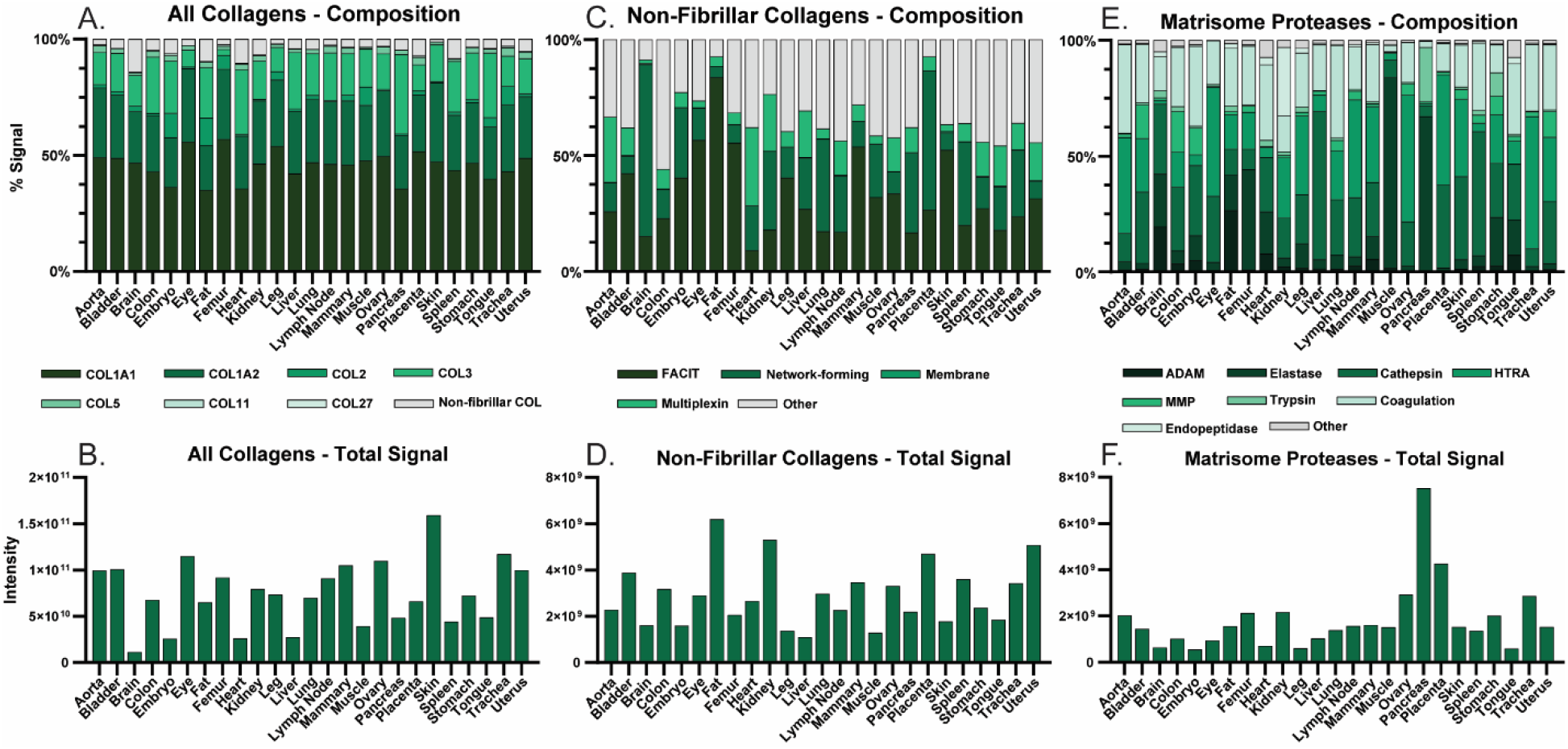
Fractional composition and abundance of ECM protein subclasses vary by tissue. A) Fractional composition of all collagen proteins by total intensity. B) Total collagen protein intensity. C) Fractional composition of non-fibrillar collagen proteins by total intensity. D) Total non-fibrillar collagen protein intensity. E) Fractional composition of matrisome proteases by total intensity. F) Total matrisome protease intensity.

Matrisome-associated protease composition and abundance was also found to vary widely between analyzed tissues. ECM protease intensity varies 13.7-fold between the highest and lowest tissues: pancreas and embryo, respectively (Figure 4F). Matrisome protease signal identified in pancreas and muscle is >65% elastase, greater than in any other tissue, although pancreas also exhibits 5-fold more total ECM protease signal than muscle (Figure 4E,F). Total elastase abundance varies 153-fold between the lowest tissue (eye) and the highest (pancreas). Other protease classes also display large differences in fractional abundance between tissues, with the protease fraction assigned to matrix metalloproteases (MMPs) and trypsin varying 98- and 110-fold between their highest and lowest tissue values, respectively (Figure 4E).

Analysis of ECM composition by matrisome class reveals strong tissue-specific enrichment of some subclasses of structural ECM proteins, including multiplexins in heart tissue and network-forming collagens in the brain. Similar differences in fractional composition were observed during analysis of other ECM protein classes, including basement membrane proteins, elastic microfibril proteins, ECM regulators, and protease inhibitors (Supplementary Figure 1). Information regarding the structural protein composition of highly unique tissue matrices, as well as tissue-specific solubility information, can be used to better recapitulate organ matrices in cell culture and organ-on-a-chip applications.

Using PHATE, a visualization method that captures both local and global nonlinear structure using an information-geometric distance between data points, networks of related structural ECM components can be observed based on peptide expression patterns^35^. Optimization of K-means clustering resulted in seven clusters exhibiting distinct expression patterns across the analyzed tissues (Figure 5). For example, fibrillar collagens 1A1, 1A2, 3A1, 5A1, and 11A1 share a similar expression pattern across organs, driven primarily by their high expression in skin, eye, and bladder as well as their low expression in brain and embryo (largely assigned to cluster 3). Cluster 3, containing many of the ECM’s principal structural components, is the most abundant observed cluster across all tissues. Collagens 2A1, and 10A1 also share a separate expression pattern from other collagens, driven by their elevated expression in trachea and femur (cluster 7) (Figure 5B). Peptides from basement membrane proteins predominantly occupy cluster 5 due to their high expression in kidney and placenta. However, expression patterns are observed for LAMA1, LAMA2, and LAMA4 that are distinct from other basement membrane proteins, suggesting that varying laminin subtype expression contributes to tissue-specific basement membrane phenotypes (Figure 5C). LAMA1 is expressed alongside provisional matrix proteins FN1 and TNC, enriched in placenta with lowest levels in brain and muscle (cluster 1), while LAMA2 peptides are highly enriched in the pancreas (cluster 6). An additional subtype of structural matrix primarily present in bladder, colon, skin, and uterus is revealed by cluster 2, defined by enrichment of COL14A1 and LAMA4 peptides (Figure 5C).

**Figure 5.**
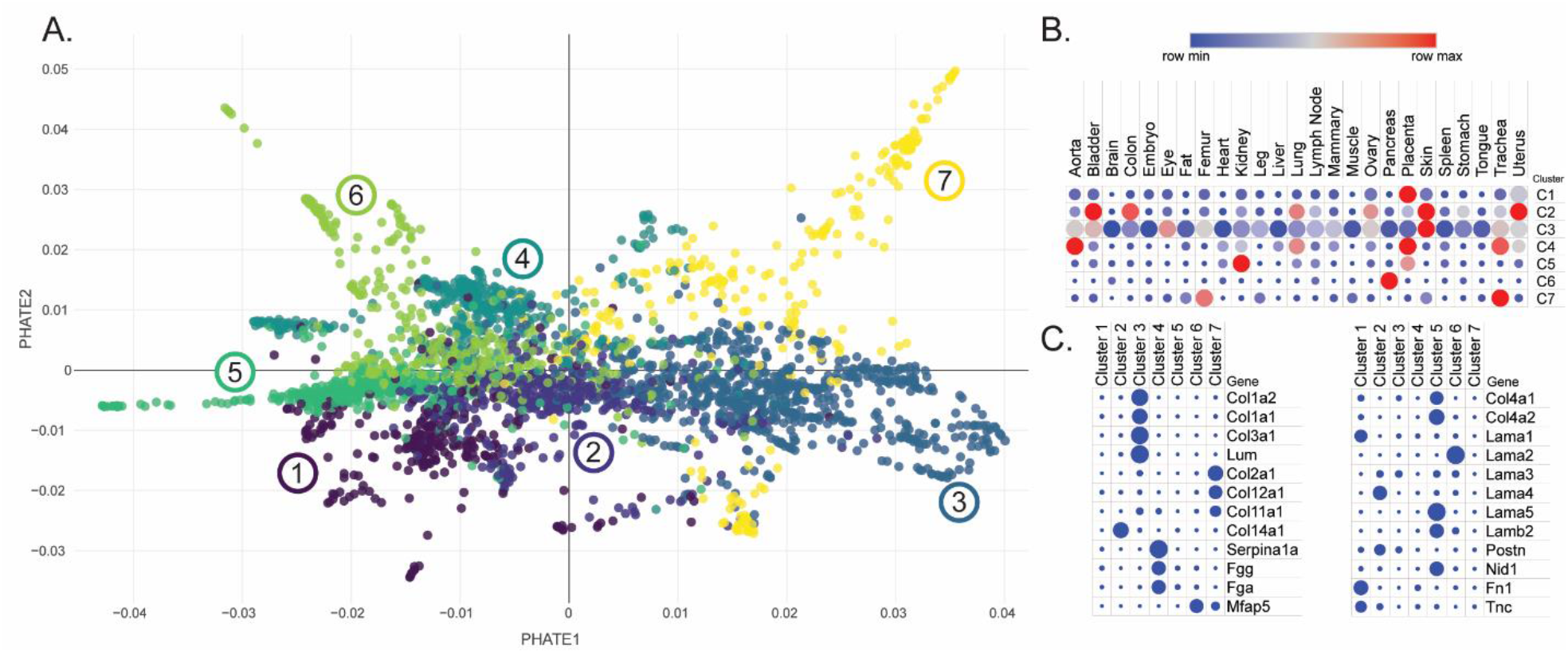
PHATE analysis of ECM peptides reveals functionally associated protein networks. A) PHATE network analysis was performed by peptide intensity and groupings were determined by K-means clustering. B) Cluster expression patterns across analyzed tissues. Red indicates row maximum intensity; blue indicates row minimum intensity. Dot size represents intensity without normalization by row. C) Representative proteins for each K-means cluster. Dot size represents number of peptides present in each cluster.

Independent networks of basement membrane proteins expressed together across analyzed tissues were also observed. Collagen 4 peptide expression is highly correlated with expression of LAMA5, B2, and C1 (suggesting laminin521), as well as NID1 and NID2, while FN1 is typically co-expressed with LAMA1 and B1 (laminin111) across tissues. More soluble ECM components, including SERPIN proteases and blood proteins such as fibrinogen, occupy K-means cluster 4, which is largely absent of peptides from structural ECM components.

Clustering of ECM peptides by expression across organs reveals both protein networks and peptide-specific expression patterns (Figure 6). Within ECM subset categories, functional protein networks can be revealed which co-express together across varying tissue matrices. For example, we observe an independent expression pattern for cartilage and connective tissue collagens COL10A1 and COL2A1. COL18A1 derived endostatin, comprised of the globular domain at the C-terminal end, is one of the most well-studied endogenous inhibitors of angiogenesis^41^. Through PHATE clustering, we observe a unique expression pattern for endostatin peptides (residues 1572-1754) compared to the remainder of the COL18A1 sequence (residues 1-1571), driven by elevated expression in bladder and stomach as well as reduced expression in lung, tongue, and colon (Figure 6B). Additionally, we observe a distinct grouping of COL12A1 peptides which cluster separately from other collagen peptides. This grouping corresponds to isoforms 3 and 5 of COL12A1, containing residues 1187-3120 of the full sequence, and has a unique expression pattern from residues 1-1186, which are contained in isoforms 1 and 2 (Figure 6C). Observed clustering of LOXL1 and LOX with ELN, suggests that enzyme-substrate expression networks can be uncovered by this form of analysis.

**Figure 6.**
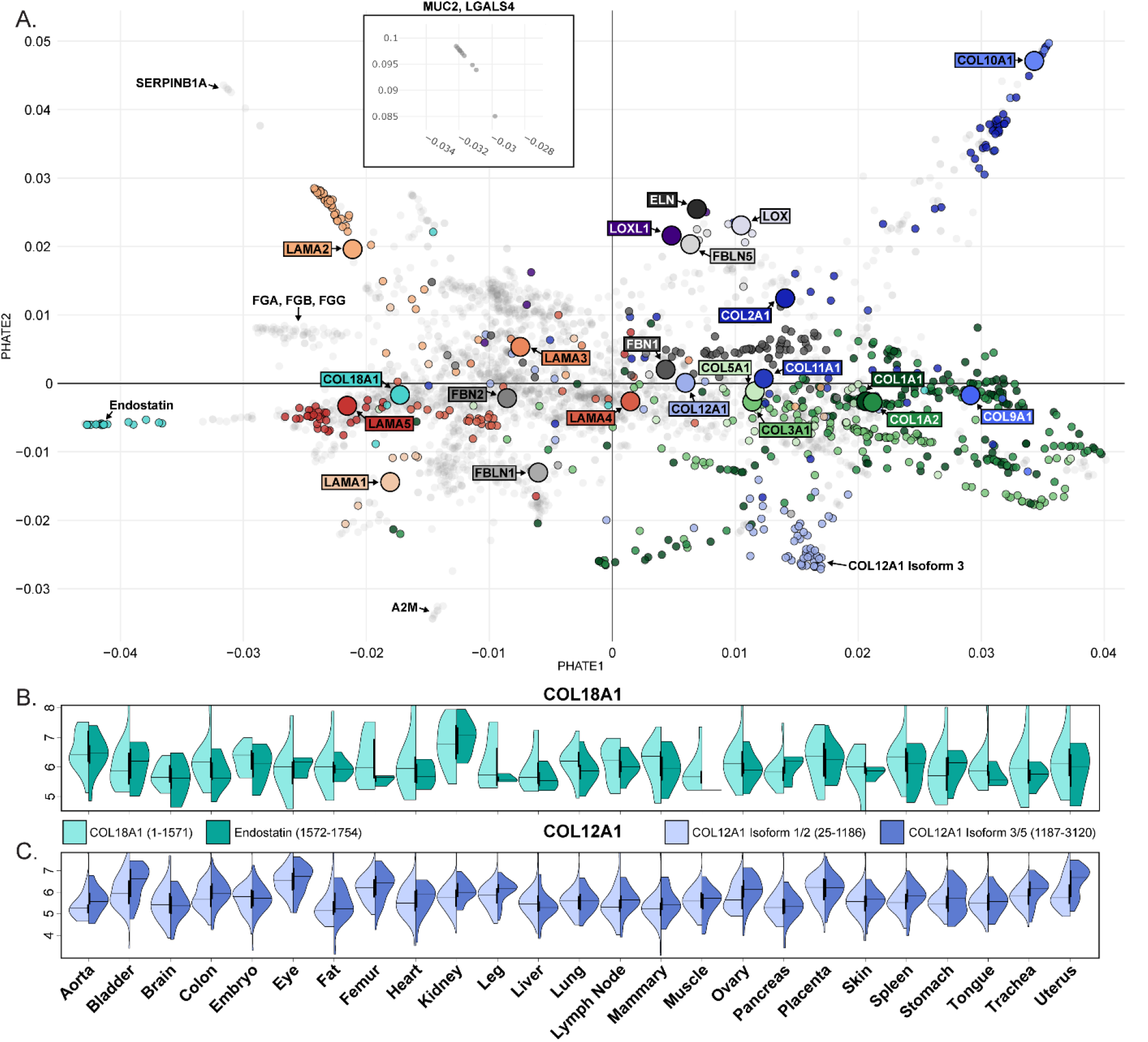
PHATE analysis of ECM peptides and proteins reveals isoform-specific expression patterns and enzyme-substrate relationships. A) PHATE network analysis performed on all ECM peptides and proteins by peptide intensity across samples. Small points represent peptides, large points represent summed protein signal. B) Split violin plot representing expression of COL18A1 residues 1-1571 (left) and residues 1572-1754 (endostatin, right) across analyzed tissues. C) Split violin plot representing expression of COL12A1 residues 1-1571 (Isoforms 1/2, left) and residues 1572-1754 (Isoforms 3/5, right) across analyzed tissues.

## Conclusions

Here we have provided ECM-optimized quantitative analysis of 25 organ proteomes. In general, collagen I is the most abundant ECM protein across tissues and a core set of ECM proteins, 34% of core ECM proteins accounting for greater than 99% AUC signal, are shared between tissues. The results suggest that tissue-specific ECM profiles are primarily determined by the ratios of these shared proteins, including collagen subtypes and other ECM components. Tissues also vary greatly in their ECM diversity, with some tissue matrices composed almost entirely of a relatively small number of ECM proteins while others contain many more ECM proteins expressed at similar abundances.

While the ECM-focused 3-step extraction method used here provides a more comprehensive ECM proteomic profile than previous tissue atlas projects, possible improvements to increase the depth of proteome coverage may be worth investigating in future studies. Despite the relatively small number of ECM components found in organs, identifications can be challenging due to the large dynamic range of protein abundances. Here we detected ECM protein abundances across 5 orders of magnitude, with COL1A1 observed as the most abundant ECM component in every tissue, whereas COL25A1 and THBS2 represent examples of proteins of low abundance found in a subset of tissues. In general, ECM proteins with more sparse identification across organs were of lower abundance. However, sample level fractionation and data-independent acquisition (DIA) approaches would enhance detection of low abundance proteins. This study was performed on a single gender and age of mouse, impeding our ability to observe gender or age specific differences or tissue variability between organisms. Hydroxylamine chemical digest has been shown to perform better than other published methods for characterization of insoluble ECM material^38^, but also induces oxidation of some amino acid residues making estimates of modification and adducts challenging.

Although our ECM-optimized method maximizes the high-confidence identifications of most highly insoluble ECM components, elastin (ELN) largely evades detection. ELN is a fundamental ECM protein essential to the elasticity of many tissues and is largely resistant to both trypsin and hydroxylamine cleavage due to extensive lysyl oxidase-based lysine crosslinking and the lack of lysine and arginine residues or hydroxylamine cleavage sites in the sequence. The low coverage of ELN in our proteomics dataset highlights the necessity to further develop protein extraction strategies that will enable accurate identification and quantification of all ECM proteins.

By defining the tissue-specific protein composition of a wide variety of matrices, we can begin to evaluate the relationship between protein components, cellular responses, and tissue biomechanical properties. Understanding the interaction between composition and biomechanics will allow for better tuning of engineered matrices for use in 3D culture and regenerative medicine applications. Due to complex interactions between cells and a wide variety of components in the surrounding matrix, it has historically been difficult to engineer matrices which match the performance of native matrices for cell culture. Comprehensive characterization of matrix composition paired with correlation to application-specific outcomes will allow for matrices to be optimized for a wide variety of uses in research and medicine. The generation of an ECM atlas containing characterization of 25 mouse organs represents a significant step toward this goal.

## Data availability

The mass spectrometry proteomics data have been deposited to the ProteomeXchange Consortium via the MassIVE partner repository with the dataset identifier PXD032000.

## Acknowledgments

This work was supported by the NIH (Grant No. R33CA183685, RM1GM131968, P01HL152961, R01HL146519) and the University of Colorado Cancer Center Support Grant (P30CA046934). We would like to thank Monika Dzieciatkowska for maintaining the analytical platform used for this work.

## Notes

### Competing Interest Statement

The authors have declared no competing interest.

## References

1. Mecham, R. P. Overview of extracellular matrix. Curr. Protoc. Cell Biol. 1–16 (2012). doi:10.1002/0471143030.cb1001s57

2. Hynes, R. O. Extracellular matrix: not just pretty fibrils. Science (80-.). 326, 1216–1219 (2009).

3. Yurchenco, P. D., Birk, D. E. & Mecham, R. P. Extracellular Matrix Assembly and Structure. (Academic press, 1994).

4. Bonnans, C., Chou, J. & Werb, Z. Remodelling the extracellular matrix in development and disease. Nat Rev Mol Cell Biol. 15, 786–801 (2014).

5. Smith, M. M. & Melrose, J. Proteoglycans in Normal and Healing Skin. Adv. Wound Care 4, 152–173 (2015).

6. Szauter, K. M., Cao, T., Boyd, C. D. & Csiszar, K. Lysyl oxidase in development, aging and pathologies of the skin. Pathol. Biol. 53, 448–456 (2005).

7. Barrett, A. S., Maller, O., Pickup, M. W., Weaver, V. M. & Hansen, K. C. Compartment resolved proteomics reveals a dynamic matrisome in a biomechanically driven model of pancreatic ductal adenocarcinoma. J. Immunol. Regen. Med. 1, 67–75 (2018).

8. Wynn, T. a. Cellular and molecular mechanisms of fibrosis. J Pathol 46, 26–32 (2008).

9. Uhlen, M. et al. Tissue-based map of the human proteome. Science (80-.). 347, (2015).

10. Kim, M.-S. et al. A draft map of the human proteome. Nature 509, 575–581 (2014).

11. Wilhelm, M. et al. Mass-spectrometry-based draft of the human proteome. Nature 509, 582–587 (2014).

12. Frantz, C., Stewart, K. M. & Weaver, V. M. The extracellular matrix at a glance. J. Cell Sci. 123, 4195–4200 (2010).

13. Hill, R. C., Calle, E. A., Dzieciatkowska, M., Niklason, L. E. & Hansen, K. C. Quantification of Extracellular Matrix Proteins from a Rat Lung Scaffold to Provide a Molecular Readout for Tissue Engineering. Mol. Cell. Proteomics 14, 961–973 (2015).

14. Schmidt, T. et al. ProteomicsDB. Nucleic Acids Res. 46, D1271–D1281 (2018).

15. Mora Huertas, A. C., Schmelzer, C. E. H., Hoehenwarter, W., Heyroth, F. & Heinz, A. Molecular-level insights into aging processes of skin elastin. Biochimie 128–129, 163– 173 (2016).

16. Wang, M. et al. PaxDb, a Database of Protein Abundance Averages Across All Three Domains of Life. Mol. Cell. Proteomics 11, 492–500 (2012).

17. Abbas, M. et al. Vertebrate cell culture as an experimental approach – limitations and solutions. Comp. Biochem. Physiol. Part - B Biochem. Mol. Biol. 254, 110570 (2021).

18. Edmondson, R., Broglie, J. J., Adcock, A. F. & Yang, L. Three-dimensional cell culture systems and their applications in drug discovery and cell-based biosensors. Assay Drug Dev. Technol. 12, 207–218 (2014).

19. Sato, T. et al. Single Lgr5 stem cells build crypt-villus structures in vitro without a mesenchymal niche. Nature 459, 262–265 (2009).

20. Barrila, J. et al. Modeling host-pathogen interactions in the context of the microenvironment: Three-dimensional cell culture comes of age. Infect. Immun. 86, (2018).

21. Blondel, D. & Lutolf, M. P. Bioinspired hydrogels for 3d organoid culture. Chimia (Aarau). 73, 81–85 (2019).

22. Clevers, H. Modeling Development and Disease with Organoids. Cell 165, 1586–1597 (2016).

23. Chaudhuri, O. Viscoelastic hydrogels for 3D cell culture. Biomater. Sci. 5, 1480–1490 (2017).

24. Nehrenheim, L. et al. Native aortic valve derived extracellular matrix hydrogel for three dimensional culture analyses with improved biomimetic properties. Biomed. Mater. 14, (2019).

25. Nurmenniemi, S. et al. A novel organotypic model mimics the tumor microenvironment. Am. J. Pathol. 175, 1281–1291 (2009).

26. Hansen, K. C. et al. An In-solution Ultrasonication-assisted Digestion Method for Improved Extracellular Matrix Proteome Coverage. Mol. Cell. Proteomics 8, 1648–1657 (2009).

27. Laklai, H. et al. Genotype tunes pancreatic ductal adenocarcinoma tissue tension to induce matricellular-fibrosis and tumor progression. Nat Med. 22, 497–505 (2016).

28. Goddard, E. T. et al. Quantitative extracellular matrix proteomics to study mammary and liver tissue microenvironments. Int J Biochem Cell Biol 81, 223–232 (2016).

29. Goddard, E. T. et al. The rodent liver undergoes weaning-induced involution and supports breast cancer metastasis. Cancer Discov. 7, 177–187 (2017).

30. Hill, R. C. et al. Preserved Proteins from Extinct Bison latifrons Identified by Tandem Mass Spectrometry; Hydroxylysine Glycosides are a Common Feature of Ancient Collagen. Mol. Cell. Proteomics 14, 1946–1958 (2015).

31. Barrett, A. S. et al. Hydroxylamine chemical digestion for insoluble extracellular matrix characterization. J Proteome Res. 44, 319–335 (2017).

32. Wiśniewski, J. R., Zougman, A., Nagaraj, N. & Mann, M. Universal sample preparation method for proteome analysis. Nat. Methods 6, 359–362 (2009).

33. Mellacheruvu, D. et al. The CRAPome: A contaminant repository for affinity purification-mass spectrometry data. Nat. Methods 10, 730–736 (2013).

34. Shao, X., Taha, I. N., Clauser, K. R., Gao, Y. (Tom) & Naba, A. MatrisomeDB: The ECM-protein knowledge database. Nucleic Acids Res. 48, D1136–D1144 (2020).

35. Moon, K. R. et al. Visualizing structure and transitions in high-dimensional biological data. Nat. Biotechnol. 37, 1482–1492 (2019).

36. Wang, S. et al. Integrated view and comparative analysis of baseline protein expression in mouse and rat tissues. bioRxiv (2021).

37. Uhlén, M. et al. Tissue-based map of the human proteome. Science (80-.). 347, (2015).

38. McCabe, M. C. et al. Evaluation and Refinement of Sample Preparation Methods for Extracellular Matrix Proteome Coverage. Mol. Cell. Proteomics 20, (2021).

39. McCabe, M. C. et al. Alterations in Extracellular Matrix Composition during Aging and Photoaging of the Skin. Matrix Biol. Plus (2020).

40. Schiller, H. B. et al. Time- and compartment-resolved proteome profiling of the extracellular niche in lung injury and repair. Mol. Syst. Biol. 11, 1–19 (2015).

41. Walia, A. et al. Endostatin’s Emerging Roles in Angiogenesis, Lymphangiogenesis, Disease, and Clinical Applications. Biochim Biophys Acta. 1850, 2422–2388 (2015).

