## Supplementary Information for "A mass spectrometry-based atlas of extracellular matrix proteins across 25 mouse organs"

| | |
|---|---|
| Collagens | Col12a1, Col14a1, Col16a1, Col9a1, Col9a2, Col11a1, Col11a2, Col1a1, Col1a2, Col27a1, Col2a1, Col3a1, Col5a1, Col5a2, Col17a1, Col25a1, Col10a1, Col8a1, Col8a2, Col26a1, Col28a1, Col6a1, Col6a2, Col6a4, Col6a5, Col6a6, Col7a1, Col15a1, Col18a1, Col4a1, Col4a2, Col4a3, Col4a4 |
| Non-Fibrillar Collagens | Col14a1, Col16a1, Col9a1, Col9a2, Col17a1, Col25a1, Col10a1, Col8a1, Col8a2, Col26a1, Col28a1, Col6a1, Col6a2, Col6a4, Col6a5, Col6a6, Col7a1, Col15a1, Col18a1, Col4a1, Col4a2, Col4a3, Col4a4 |
| ECM Proteases | Cpn2, Habp2, Adam10, Adam11, Adam17, Adam22, Adam23, Adam9, Adamts20, Adamts5, Adamtsl4, Ctsa, Ctsb, Ctsc, Ctsd, Ctse, Ctsf, Ctsg, Ctsh, Ctsj, Ctsl, Ctss, Ctsz, F10, F12, F2, F7, F9, Plg, Masp1, Masp2, Cela1, Cela2a, Cela3b, Elane, Mep1a, Mep1b, Htra1, Htra3, Mmp12, Mmp13, Mmp2, Mmp9, Prss2 |
| Basement Membrane | Col15a1, Col18a1, Col4a1, Col4a2, Col4a3, Col4a4, Agrn, Lama1, Lama2, Lama3, Lama4, Lama5, Lamb1, Lamb2, Lamb3, Lamc1, Lamc2, Lamc3, Nid1, Nid2, Hspg2 |
| ECM Regulators | Plod1, Plod2, Plod3, Hpse2, Hyal2, Sulf2, Pzp, St14, Cpn2, Habp2, Cd109, Hrg, Kng1, Slpi, Adam10, Adam11, Adam17, Adam22, Adam23, Adam9, Adamts20, Adamts5, Adamtsl4, Ctsa, Ctsb, Ctsc, Ctsd, Ctse, Ctsf, Ctsg, Ctsh, Ctsj, Ctsl, Ctss, Ctsz, F13a1, F10, F12, F2, F7, F9, Plg, Masp1, Masp2, Cst10, Cst3, Cstb, Ngly1, Cela1, Cela2a, Cela3b, Elane, Mep1a, Mep1b, Htra1, Htra3, Itih1, Itih2, Itih3, Itih4, Itih5, Fam20b, Fam20c, Lox, Loxl1, Loxl2, Loxl3, Loxl4, A2m, Ambp, Mug2, Mmp12, Mmp13, Mmp2, Mmp9, Ogfod1, P4ha1, P4ha2, Agt, Serpina10, Serpina12, Serpina1a, Serpina1b, Serpina1d, Serpina1e, Serpina3b, Serpina3k, Serpina3m, Serpina3n, Serpina6, Serpinb12, Serpinb1a, Serpinb1b, Serpinb2, Serpinb5, Serpinb7, Serpinb8, Serpinc1, Serpind1, Serpine1, Serpine2, Serpinf1, Serpinf2, Serping1, Serpinh1, Serpini2, Tgm1, Tgm2, Tgm3, Tgm5, Timp2, Timp3, Prss2 |
| Elastic Microfibrils | Eln, Emilin1, Emilin2, Emilin3, Fbln1, Fbln2, Fbln5, Fbn1, Fbn2, Mfap1a, Mfap2, Mfap4, Mfap5, Vtn |
| ECM Protease Inhibitors | Papln, Cd109, Hrg, Kng1, Slpi, Cst10, Cst3, Cstb, Itih1, Itih2, Itih3, Itih4, Itih5, A2m, Ambp, Mug2, Agt, Serpina10, Serpina12, Serpina1a, Serpina1b, Serpina1d, Serpina1e, Serpina3b, Serpina3k, Serpina3m, Serpina3n, Serpina6, Serpinb12, Serpinb1a, Serpinb1b, Serpinb2, Serpinb5, Serpinb7, Serpinb8, Serpinc1, Serpind1, Serpine1, Serpine2, Serpinf1, Serpinf2, Serping1, Serpinh1, Serpini2, Timp2, Timp3 |

**Supplementary Table 1.** Contents of matrisome subcategories used in ECM composition analysis.

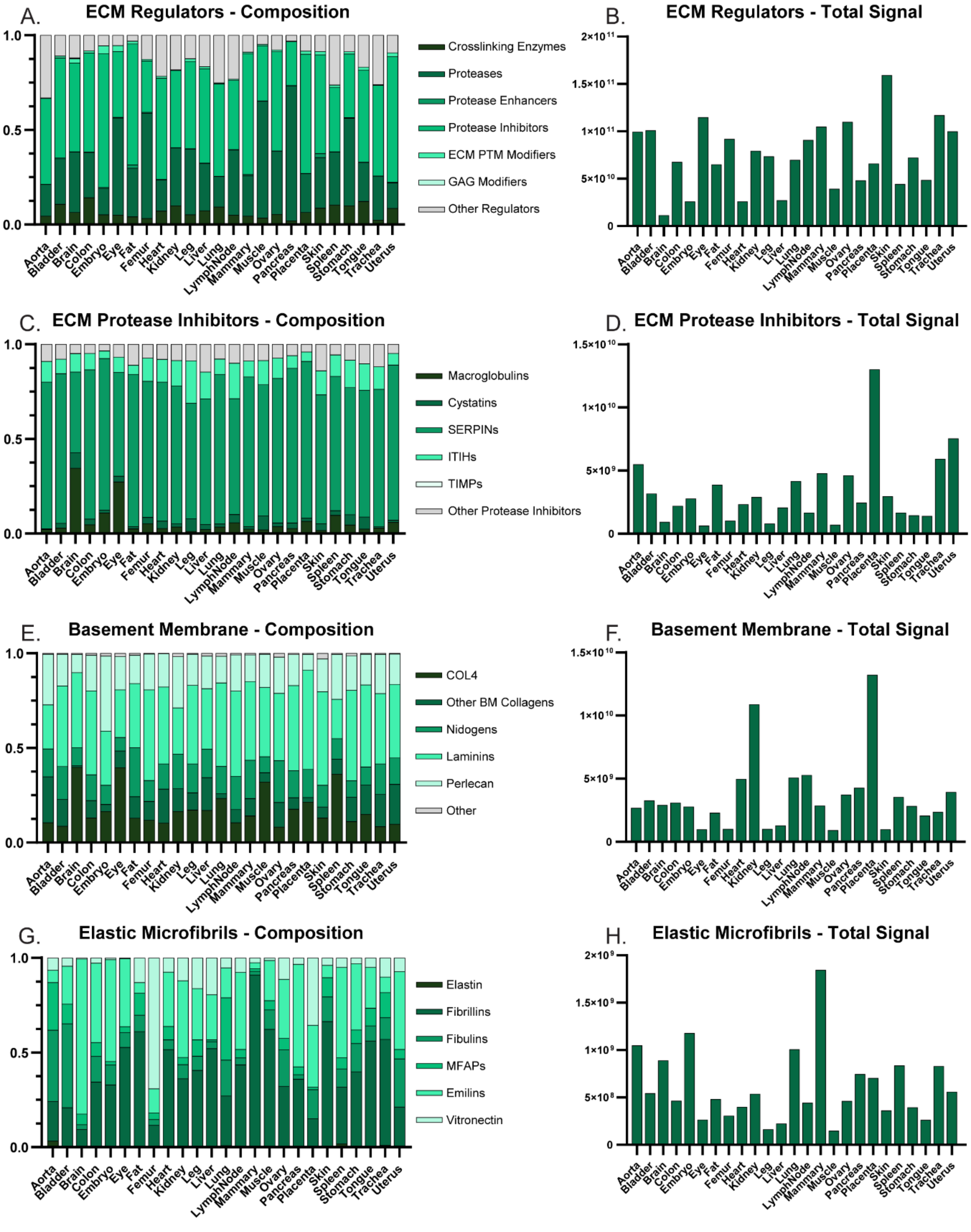


**Supplementary Figure 1. Fractional composition and abundance of additional ECM protein subclasses vary by tissue.** A) Fractional composition of ECM regulators by total intensity. B) Total ECM regulator protein intensity. C) Fractional composition of ECM protease inhibitors by total intensity. D) Total ECM protease inhibitor intensity. E) Fractional composition of basement membrane proteins by total intensity. F) Total basement membrane protein intensity. G) Fractional composition of elastic microfibril proteins by total intensity. H) Total elastic microfibril protein intensity.

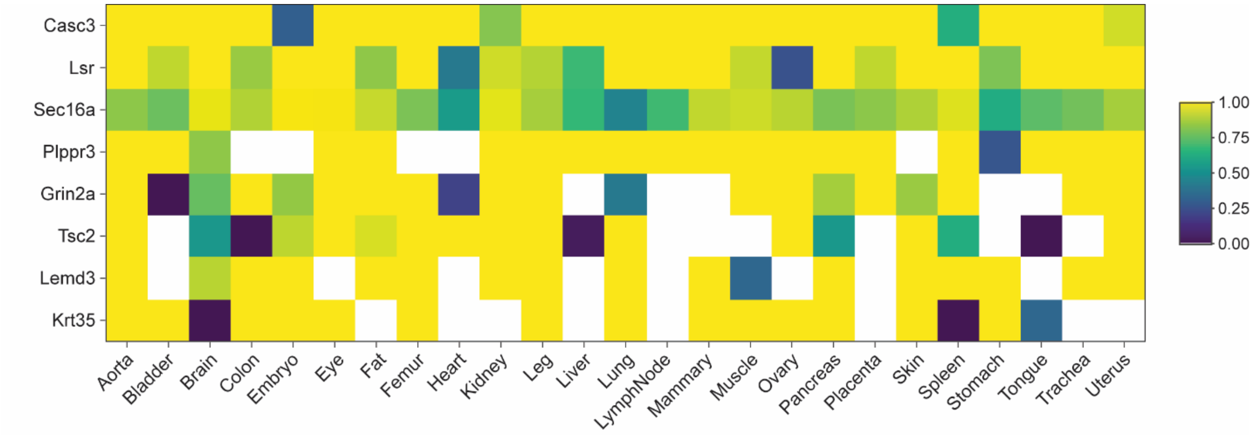


**Supplementary Figure 2. Cellular proteins resistant to chaotrope extraction.** Cells are colored by solubility, calculated as the percent of total intensity for each protein identified in the iECM fraction.
